# Disparate effects of metformin on *Mycobacterium tuberculosis* infection in diabetic and non-diabetic mice

**DOI:** 10.1101/2020.07.17.209734

**Authors:** Harindra D. Sathkumara, Karyna Hansen, Socorro Miranda-Hernandez, Brenda Govan, Catherine M. Rush, Lars Henning, Natkunam Ketheesan, Andreas Kupz

**Author notes:** Corresponding Author Address correspondence to Dr Andreas Kupz.

## Abstract

Comorbid type 2 diabetes poses a great challenge to the global control of tuberculosis. Here we assessed the efficacy of metformin (MET); an anti-diabetic drug, in mice infected with a very-low dose of *Mycobacterium tuberculosis*. In contrast to diabetic mice, infected non-diabetic mice that received the same therapeutic concentration of MET presented with significantly higher disease burden. This warrants further studies to investigate the disparate efficacy of MET against tuberculosis in diabetic and non-diabetic individuals.

## Introduction

Tuberculosis (TB) remains one of the deadliest infectious diseases with an estimated annual mortality of 1.5 million and nearly 1.7 billion latently-infected people worldwide.^1^ Whilst infections with drug-susceptible *Mycobacterium tuberculosis* (*Mtb*), the causative agent of TB, can be treated with long-term antibiotic therapy, emergence of drug-resistant strains and increasing incidence of comorbid conditions, such as diabetes mellitus (DM), pose a great challenge to TB eradication.^2^ It is estimated that 463 million people are currently living with diabetes^3^ and have a threefold increased risk of developing active TB^4^ and demonstrated a strong link with multi-drug-resistant (MDR) TB.^5^

Poor treatment adherence, clinical complications and continuous exposure to conventional anti-TB monotherapy often lead to drug tolerance and resistance.^6^ In addition to the evaluation of new and existing repurposed anti-TB drugs, there has also been an increased interest in non-antimicrobial host-directed therapies (HDTs) which often target host immune responses with the potential to shorten and improve treatment duration and therapeutic efficacy against all forms of TB.^7^

Metformin (MET; 1,1-dimethylbiguanide) is a widely prescribed AMP-activated protein kinase (AMPK)-activating anti-diabetic drug found to be associated with reduced TB risk among diabetic patients.^8, 9^ MET was previously shown to inhibit intracellular growth of *Mtb* and improve the efficacy of first-line anti-TB drug; isoniazid (INH) in young non-diabetic C57BL/6 mice.^9^ However, in a recent experiment, MET failed to enhance the potency of current anti-TB treatment regimen in young BALB/c mice^10^, implying the need to resolve the discrepant findings between experiments and more importantly to investigate the true impact of MET on TB in the context of diabetes.

In this study, we sought to simultaneously evaluate the protective efficacy of MET alone and in combination with INH against TB using a robust model of murine T2D and age-matched non-diabetic control mice.^11^

## Materials and methods

### Ethics

This study was conducted in accordance with the National Health and Medical Research Council (NHMRC) animal care guidelines with all procedures approved by the animal ethics committee (A2400) of James Cook University (JCU), Australia.

### Diet-induced murine T2D

C57BL/6 male mice housed under specific-pathogen free (SPF) conditions were randomly divided into two groups for dietary interventions. One group was given *ad libitium* access to an energy dense diet (EDD; 23% fat, 19% protein, 50.5% dextrose and 7.5% fibre), while the second group received an isometric quantity of standard rodent diet (SD). Thirty weeks following diet intervention, mice were assessed for body weight, fasting blood glucose levels and glucose tolerance.^11^ Area under the curve (AUC) calculated from glucose tolerance test (GTT) was used to confirm the status of diabetes.^11^

### Bacteria

*Mtb* H37Rv were cultured in 10% ADC enriched Middlebrook 7H9 broth (BD Biosciences) supplemented with 0.2% glycerol and 0.05% Tween 80. Mid-logarithmic cultures were harvested, washed in sterile PBS and stored in 15% glycerol/PBS at -80°C.

### Aerosol infection

Mice were infected with a very-low aerosol dose (10-20 CFUs) of *Mtb* H37Rv using a Glas-Col inhalation exposure system in a biosafety level 3 (BSL3) laboratory. The infectious dose was determined by culturing homogenized lung tissues of 4-5 mice on 10% OADC enriched 7H11 agar plates 1-day post infection (p.i.).

### Drugs

Seven days post aerosol *Mtb* challenge, treatments were initiated by administering MET and INH (both from Sigma) alone or in combination (MET+INH) to mice in drinking water delivering ∼500 mg/kg and ∼10 mg/kg, respectively (1.25 mg/ml MET in drinking water delivers a dose of approximately 250 mg/kg to mice^12^). Water bottles containing drugs were changed every 4-6 days.

### Sample collection

At designated time points, mice were sacrificed and blood was collected in Z-gel tubes (Sarstedt). Coagulated blood was spun and serum samples were filtered using 0.2 µm SpinX columns (Sigma) for storage at -20^°^C. Lungs were aseptically removed for CFU enumeration and histology analysis.

### CFU enumeration

Right lung lobes were homogenized in sampling bags containing 1 ml of sterile PBS/ 0.05% Tween 80. Serial dilutions of tissue homogenates were plated on 10% OADC enriched 7H11 agar plates supplemented with 10 µg/ul cycloheximide and 20 µg/ml ampicillin. Plates were incubated for 3-4 weeks at 37^°^C. Total CFUs per each organ was calculated based on dilution factor and organ size.

### Lung histology

Left lung lobes were fixed overnight with 10% neutral buffered formalin and transferred into 70% ethanol the next day. Processed lung lobes were embedded in paraffin, cut (4 µm sections) and stained with Hematoxylin and Eosin (H&E) and Ziehl-Neelsen (ZN). ImageJ software (NIH) was used to measure the total surface area followed by the areas of dense cell infiltration. Proportions of infiltration were calculated accordingly.

### Serum cytokine/chemokine analysis

Serum samples were prepared for Bio-Plex Pro Mouse Cytokine 23-Plex assay (BioRad) following manufacturer’s specifications. Measurements were taken using MagPix (Luminex) instrument. Log_2_ concentration of each analyte is visualized in a heat map.

### Study inclusion

Mice which showed both CFUs on 7H11 agar plates and *Mtb* bacilli by ZN stain were included. Due to the absence of viable CFUs on agar plates 5 mice were excluded from analysis despite confirmed presence of *Mtb* using ZN staining and increased lung inflammation at 45 days p.i. **(Figure S1-2)**.

### Data Analysis

Statistical analysis was performed using GraphPad Prism version 8. Two or multiple group analysis was carried out using Student’s *t*-test and one-way analysis of variance (ANOVA), followed by Dunnett’s multiple comparison test, respectively. *P<*0.05 was considered significant.

## Results

### Murine model of T2D

Following 30 weeks of diet intervention **(Figure 1a)**, mice fed with EDD presented with a significantly increased body weight **(Figure 1b)**, elevated fasting blood glucose levels **(Figure 1c)** and impaired glucose tolerance as reflected by AUC **(Figure 1d)**; hall mark features of human T2D.^11^ Confirmed T2D and non-diabetic control mice were subjected to downstream experimental procedures.

**Figure 1:**
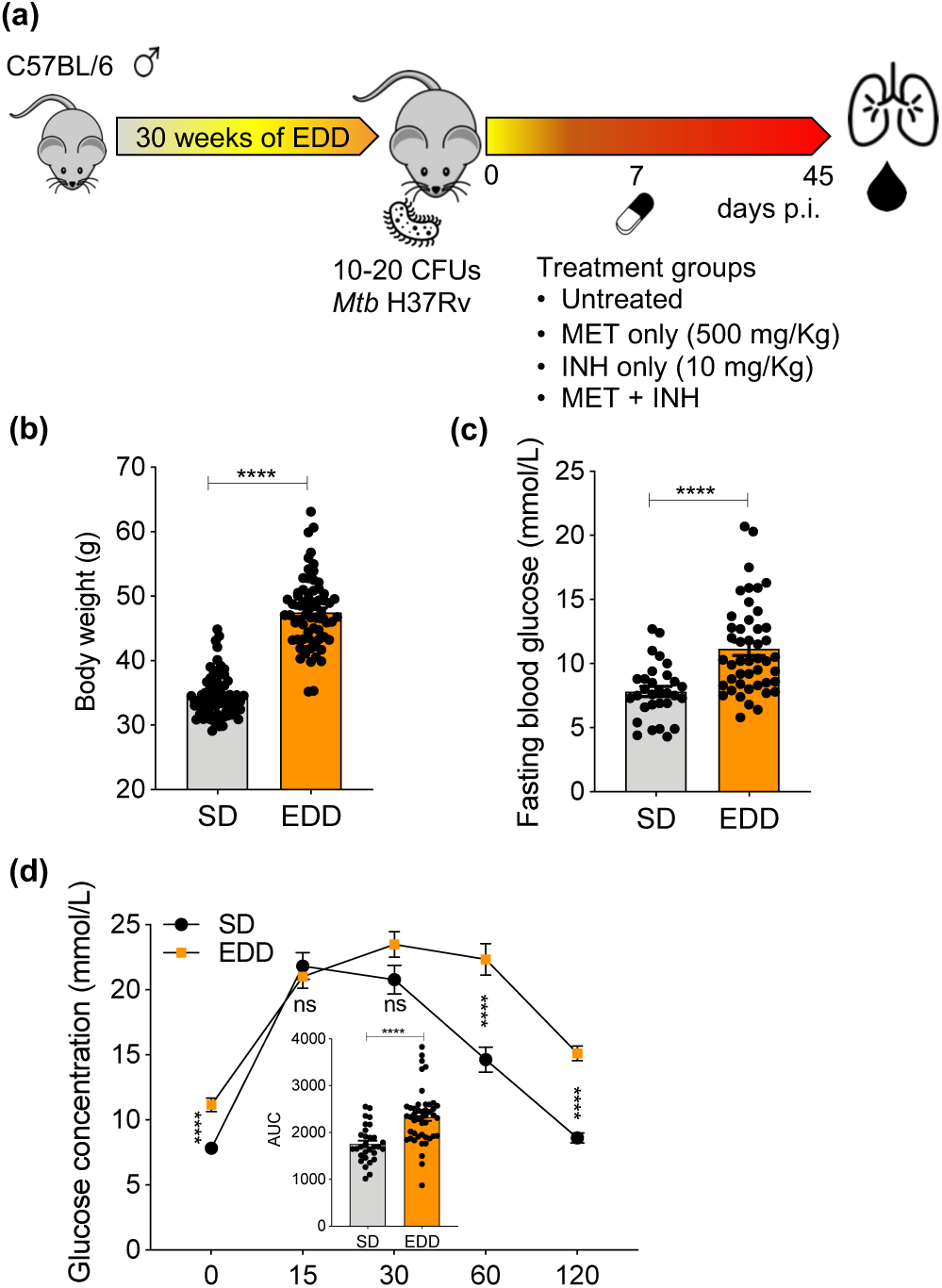
Diet induced model of murine T2D, *Mtb* infection and treatments. **(a)** Six to eight weeks old C57BL/6 male mice were fed with EDD and SD (control mice) for 30 weeks to induce murine T2D. Following dietary intervention mice were assessed for **(b)** body weight, **(c)** fasting blood glucose levels and **(d)** glucose tolerance. **(d)** GTT was performed by measuring glucose concentrations at 15, 30, 60, and 120 mins post i.p. glucose administration (2 g/kg) and calculating AUC. **(a)** Diabetic and non-diabetic control mice were infected with very-low aerosol dose (10-20 CFUs) of *Mtb* H37Rv. Seven days p.i., MET (500 mg/kg), INH (10 mg/kg) and combination of MET + INH were administered in drinking water. Results are presented as individual data points **(b-d)** and pooled data means ± SEM **(d)** from two pooled independent experiments. Statistical analysis: \**p* < 0.05; \*\**p* < 0.01; \*\*\**p* < 0.001; \*\*\*\**p* < 0.0001 by Student’s *t-*test. Abbreviations: p.i.; post infection, EDD; energy dense diet, SD; standard diet, i.p.; intraperitoneal, GTT; glucose tolerance test, AUC; area under the curve.

### Divergent effects of metformin

To investigate whether metformin could restrict the growth of *Mtb* and enhance the efficacy of the first-line anti-TB drug INH, we infected non-diabetic control and T2D mice with a very-low aerosol dose of *Mtb* H37Rv which closely mimics the human TB. CFUs recovered from lung tissue 7 days p.i. revealed that both control and T2D mice carried a similar bacterial burden prior to treatment **(Figure 2a)**. At 45 days p.i., 10 mg/kg INH sterilized *Mtb* infection in almost all animals **(Figure 2b)** thus, combination therapy with 500 mg/kg MET did not further enhance the efficacy of INH in our experimental settings **(Figure 2b)**. Both INH and MET+INH treatments were also accompanied by significantly lower lung immunopathology **(Figure 2c** and **S1)**. Interestingly, MET-treated T2D mice had a significantly lower overall bacterial burden (∼1.5-log reduction compared to untreated T2D; **Figure 2b)**. Strikingly, in contrast to a previous report,^9^ non-diabetic control mice treated with MET demonstrated a substantially increased lung *Mtb* burden (**Figure 2b)** and unchanged pro-inflammatory IL-12p40 levels **(Figure 2d)** reflecting an ongoing inflammation. This disparate efficacy of MET in non-diabetic and T2D mice resulted in a ∼2-log difference in lung *Mtb* burden. Overall, systemic cytokine/ chemokine levels were comparable between control and T2D mice **(Figure S3a)** and all treatments seem to have downregulated a majority of analytes **(Figure S3b)**.

**Figure 2:**
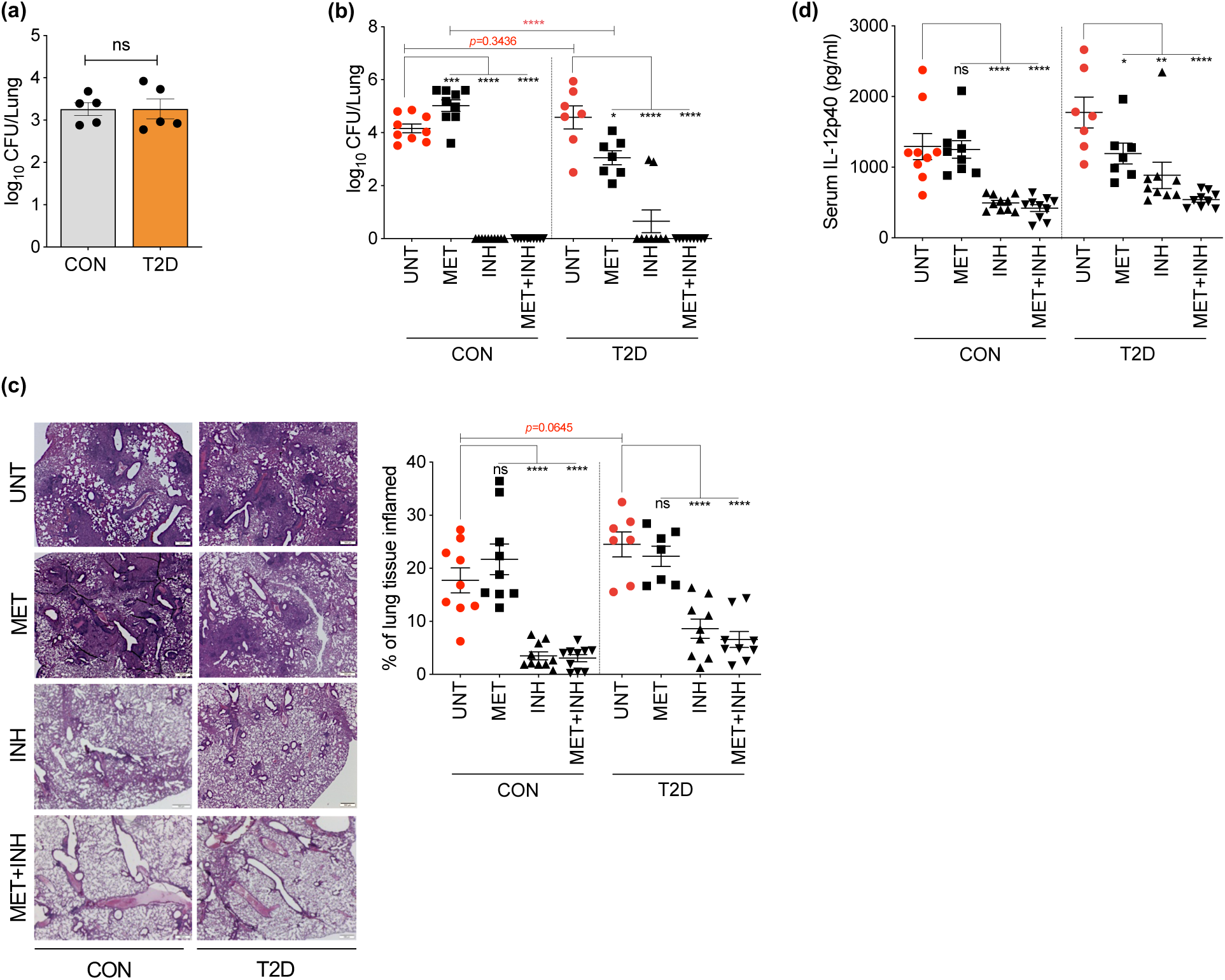
Divergent effects of MET on control and T2D mice. Seven days p.i., group of mice were sacrificed and assessed for **(a)** lung bacterial loads. Forty-five days following infection, treated and untreated mice from both control and T2D groups were assessed for **(b)** viable bacteria in lung, (**c)** lung inflammation and **(d)** serum IL-12p40 levels. Results are presented as individual data points **(a-d)** and representative images (25x) **(c)** from two pooled independent experiments (n=7-10 mice per group). Statistical analysis: \**p* < 0.05; \*\**p* < 0.01; \*\*\**p* < 0.001; \*\*\*\**p* < 0.0001 by Student’s *t*-test **(b, c)** and One-way ANOVA followed by Dunnett’s multiple comparison test **(b-d)** from 7-10 mice per group from two pooled independent experiments. Abbreviations: CON; control, T2D; type 2 diabetes, CFU; colony forming units, p.i.; post infection, UNT; untreated (Solid red circles), MET; metformin (solid black squares), INH; Isoniazid (solid black up-pointing triangles), MET+INH; combined therapy (solid black down-pointing triangles).

## Discussion

Cumulative evidence suggests that MET prescription is associated with a significantly lower incidence of active TB among TB/DM comorbid patients.^13^ Reduced mortality, fewer pulmonary cavities and rapid culture conversion are evidence of improved health outcome. In addition, Singhal and colleagues reported that MET treatment was also associated with reduced LTBI prevalence and enhanced *Mtb*-specific T cell responses as determined by T-SPOT assay.^9^ However, no significant association between LTBI and TB/DM patients taking MET was found in a more recent study.^14^ Collectively, these retrospective studies indicate that MET has the propensity to improve the overall treatment outcome when used with existing anti-TB regimens in TB/DM comorbid patients. Conformably, therapeutic concentration of MET significantly improved the disease outcome in our T2D mice. However, MET did not enhance the sterilizing effect of INH. These discrepant findings could be due to differences in analysis time points (35 vs 45 days p.i.) and/or reduced dose of INH (5 vs 10 mg/kg) used previously.^9^ Strikingly, augmented lung bacillary loads and lung immunopathology among non-diabetic mice that received the same therapeutic MET dose indicate diminished anti-TB immunity in these mice. *Ex vivo* stimulation and *in vivo* administration of MET significantly diminished the production of *Mtb* lysate-induced pro-inflammatory cytokines and downregulated the expression of type 1 interferon pathway, respectively, in healthy human peripheral blood mononuclear cells (PBMCs).^15^

The potential role of MET as an adjunctive therapy for TB is exciting. However, current evidences for MET-induced anti-TB protection have come mostly from retrospectively evaluated studies involving TB/DM comorbid patients. A number of studies have provided evidence for MET-induced reduction in pro-inflammatory threshold via possible inhibition of mammalian target of rapamycin (mTOR)^16, 17^ and/or perturbed gut microbiota.^18-20^ Whilst this immunomodulatory effect of MET can be favorable in certain high-risk populations such as diabetics with chronic inflammation, the excessive host anti-inflammatory responses can exert negative influence on the early control of *Mtb* and bacterial dissemination as observed in our non-diabetic mice. Further pre-clinical studies are therefore warranted to decipher the potentially disparate effects of MET in diabetic and non-diabetic hosts before it gains entry into clinical trials as an adjunctive anti-TB drug. In future, our long-term T2D mouse model will enable us to investigate the efficacy and optimum therapeutic concentration of MET against TB at various stages of the development of diabetes.

## Acknowledgments

We thank Chris Wright, Lachlan Pomfrett and Serrin Rowarth for assistance with PC3 and animal house operations.

## Author Contributions

A.K. and N.K. conceived of the study. H.D.S., K.H., S.M-H., C.R. and A.K. performed experiments; H.D.S. and A.K., performed data analysis. N.K., B.G., C.R. and L.H. assisted with troubleshooting and intellectual input. H.D.S. and A.K. wrote the initial manuscript. All co-authors commented extensively on the manuscript and approved it.

## Transparency declaration

None to declare.

## References

1. WHO. Global Tuberculosis Report 2019. https://apps.who.int/iris/bitstream/handle/10665/329368/9789241565714-eng.pdf?ua=1 (19 December 2019, date last accessed).

2. Amberbir A. The challenge of worldwide tuberculosis control: and then came diabetes. Lancet Glob Health 2019; 7: e390–e1.

3. Federation ID. IDF Diabetes Atlas. http://www.diabetesatlas.org/ (27th December 2019, date last accessed).

4. Al-Rifai RH, Pearson F, Critchley JA et al. Association between diabetes mellitus and active tuberculosis: A systematic review and meta-analysis. PloS one 2017; 12: e0187967.

5. Tegegne BS, Mengesha MM, Teferra AA et al. Association between diabetes mellitus and multi-drug-resistant tuberculosis: evidence from a systematic review and meta-analysis. Syst Rev 2018; 7: 161.

6. Gillespie SH. Evolution of drug resistance in Mycobacterium tuberculosis: clinical and molecular perspective. Antimicrob Agents Chemother 2002; 46: 267–74.

7. Zumla A, Chakaya J, Centis R et al. Tuberculosis treatment and management--an update on treatment regimens, trials, new drugs, and adjunct therapies. Lancet Respir Med 2015; 3: 220–34.

8. Lin SY, Tu HP, Lu PL et al. Metformin is associated with a lower risk of active tuberculosis in patients with type 2 diabetes. Respirology 2018; 23: 1063–73.

9. Singhal A, Jie L, Kumar P et al. Metformin as adjunct antituberculosis therapy. Sci Transl Med 2014; 6: 263ra159.

10. Dutta NK, Pinn ML, Karakousis PC. Metformin Adjunctive Therapy Does Not Improve the Sterilizing Activity of the First-Line Antitubercular Regimen in Mice. Antimicrob Agents Chemother 2017; 61.

11. Morris JL, Bridson TL, Alim MA et al. Development of a diet-induced murine model of diabetes featuring cardinal metabolic and pathophysiological abnormalities of type 2 diabetes. Biol Open 2016; 5: 1149–62.

12. Chandel NS, Avizonis D, Reczek CR et al. Are Metformin Doses Used in Murine Cancer Models Clinically Relevant? Cell Metab 2016; 23: 569–70.

13. Zhang M, He JQ. Impacts of metformin on tuberculosis incidence and clinical outcomes in patients with diabetes: a systematic review and meta-analysis. Eur J Clin Pharmacol 2019.

14. Magee MJ, Salindri AD, Kornfeld H et al. Reduced prevalence of latent tuberculosis infection in diabetes patients using metformin and statins. The European respiratory journal 2019; 53.

15. Lachmandas E, Eckold C, Bohme J et al. Metformin Alters Human Host Responses to Mycobacterium tuberculosis in Healthy Subjects. The Journal of infectious diseases 2019; 220: 139–50.

16. Arts RJW, Carvalho A, La Rocca C et al. Immunometabolic Pathways in BCG-Induced Trained Immunity. Cell reports 2016; 17: 2562–71.

17. Lachmandas E, Beigier-Bompadre M, Cheng SC et al. Rewiring cellular metabolism via the AKT/mTOR pathway contributes to host defence against Mycobacterium tuberculosis in human and murine cells. European journal of immunology 2016; 46: 2574–86.

18. Elbere I, Kalnina I, Silamikelis I et al. Association of metformin administration with gut microbiome dysbiosis in healthy volunteers. PloS one 2018; 13: e0204317.

19. Ma W, Chen J, Meng Y et al. Metformin Alters Gut Microbiota of Healthy Mice: Implication for Its Potential Role in Gut Microbiota Homeostasis. Front Microbiol 2018; 9: 1336.

20. Dumas A, Corral D, Colom A et al. The Host Microbiota Contributes to Early Protection Against Lung Colonization by Mycobacterium tuberculosis. Front Immunol 2018; 9: 2656.

